# Proteome-scale tagging and functional screening in mammalian cells by ORFtag

**DOI:** 10.1101/2024.01.16.575827

**Authors:** Filip Nemčko, Moritz Himmelsbach, Vincent Loubiere, Ramesh Yelagandula, Michaela Pagani, Nina Fasching, Julius Brennecke, Ulrich Elling, Alexander Stark, Stefan L. Ameres

**Author notes:** These authors contributed equally: Filip Nemčko, Moritz Himmelsbach.

## Abstract

Determining protein function in a systematic manner is a key goal of modern biology, but remains challenging with current approaches. Here, we present ORFtag, a versatile, cost-effective and highly efficient method for the massively-parallel tagging and functional interrogation of proteins at proteome scale. Using mouse embryonic stem cells, we showcase ORFtag’s utility through screens for transcriptional activators, repressors and post-transcriptional regulators. Each screen finds known and novel regulators, including long ORFs not accessible to other methods, revealing that Zfp574 is a highly selective transcriptional activator and that oncogenic fusions frequently function as transactivators.

## Main text

Proteins are pivotal in nearly all cellular processes, but their biochemical diversity often hinders systematic protein function studies. Genetic loss- or gain-of-function screens - such as CRISPR-Cas9, Cas9i and Cas9a screens - are powerful methods for identifying genes involved in specific cellular processes, but typically do not provide direct insight into protein function^1^. They are also often hampered by functional redundancies and the essentiality of many genes. Conversely, sufficiency-based assays allow direct determination of protein-inherent function^2,3^. However, current systematic methods hinge on the delivery and expression of open reading frame (ORF) libraries^4,5^, which are not only costly and challenging to maintain but also tend to favor shorter ORFs (<5kb) due to limitations in DNA synthesis, cloning, viral packaging, and delivery into cells^2^. While targeted engineering of native gene locations can bypass these limitations, recent CRISPR-Cas9 techniques for systematic gene tagging have only scaled to several hundred genes^6–10^.

Here, we present ORFtag, a versatile approach that allows for the massive, parallel, and proteome-scale tagging of endogenous ORFs, overcoming critical limitations of current methods. ORFtag is based on insertional elements such as retroviral vectors containing a constitutively active promoter, a selection gene, and a functional tag of interest followed by a splice donor sequence (**Fig. 1a**). Upon large-scale transduction of cultured cells, ORFtag cassettes randomly integrate into the genome and drive the transcription of nearby endogenous gene loci by splicing of the functional tag to splice-acceptor sites downstream of the integration site. With one cassette for each of three open reading frames, ORFtag can be used to generate N-terminal fusions of endogenous ORFs with a wide range of functional tags. A key feature of ORFtag is its compatibility with diverse functional readouts including reporter-based positive selection by fluorescence-activated cell sorting (FACS). In the selected cell population, tagged genes are identified by mapping integration sites using inverse PCR (iPCR) followed by next-generation sequencing (NGS)^11^.

**Fig. 1:**
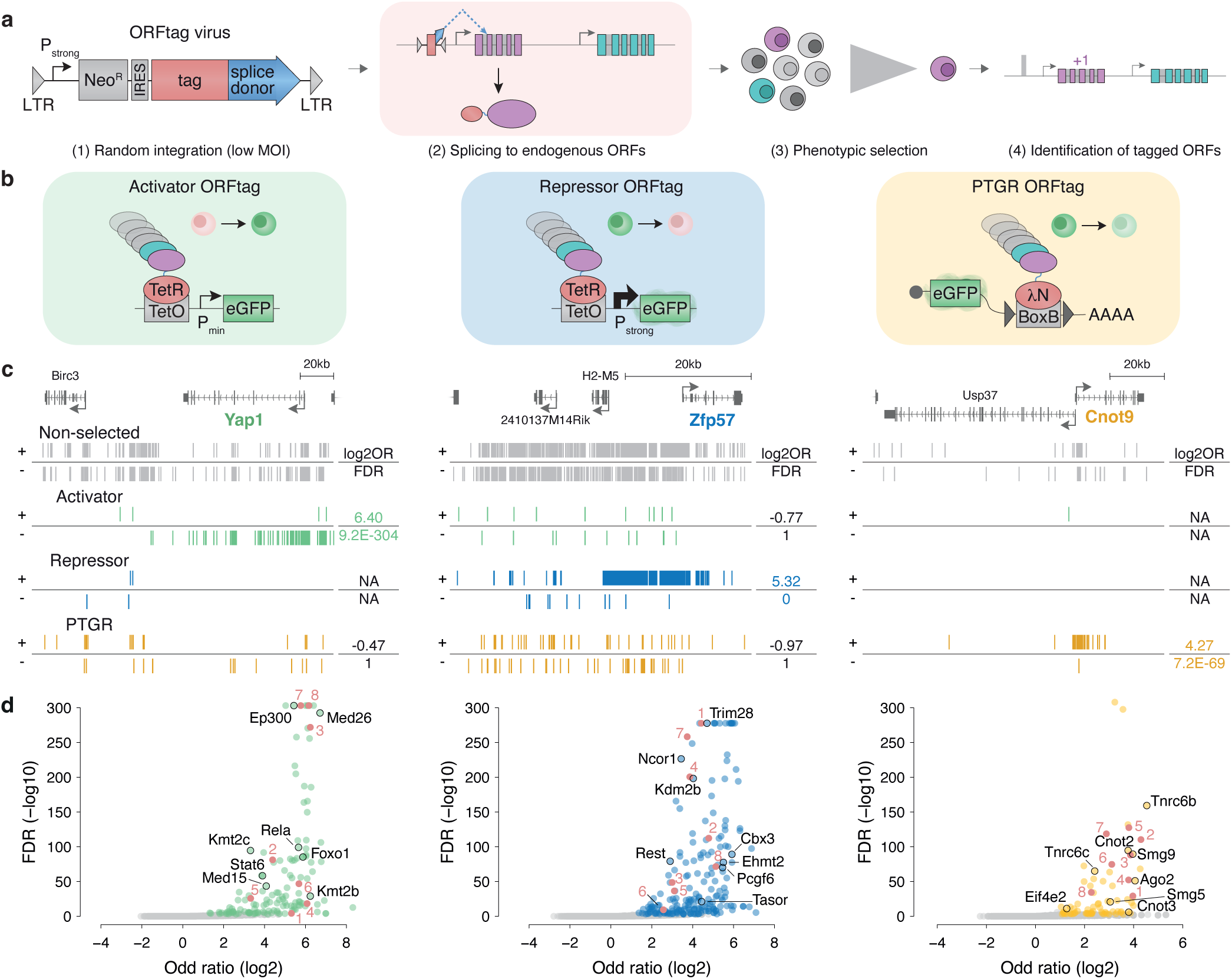
ORFtag is a versatile tool for proteome-wide functional assays. **a,** Overview of the ORFtag approach. The ORFtag cassette is embedded in a retroviral backbone and contains a constitutively active promoter, selection gene, a tag, and a splice-donor site. Upon transduction, the ORFtag cassette randomly integrates into the genome and prompts splicing to a downstream splice acceptor, ultimately producing a tagged protein. **B,** Schematic view of 3 different screens for transcriptional activators (green), repressors (blue) or PTGRs (yellow). **C,** Genome browser screenshots of ORFtag integration sites (vertical lines) in positive (+, top) or negative (-, bottom) strand direction, before (non-selected, grey) and after FACS selection at the genomic locus of each one activator (Yap1, green), repressor (Zfp57, blue) and PTGR (Cnot9, yellow) hit emerging from ORFtag screens in mESCs. Log2 odds-ratio (log2OR) and false discovery rate (FDR) are indicated. **D,** Volcano plots highlighting known (black circles, names) and validated (marked red, see Fig. 2d) hits for the 3 screens.

To benchmark the ORFtag approach, we performed three functional screens for transcriptional activators, transcriptional repressors, and post-transcriptional gene regulatory (PTGR) proteins in mouse embryonic stem cells (mESCs), each in two biological replicates (**Fig. 1b**). For the transcriptional activator and repressor screens, we systematically fused proteins to the DNA binding domain of the bacterial Tet Repressor (TetR), enabling their recruitment to TetO binding sites located upstream of an integrated GFP reporter. This reporter contained an inactive minimal promoter or a constitutively active promoter for transcriptional activator and repressor screens, respectively. For the PTGR screen, we tagged proteins with the lambda phage N protein (λN) in order to recruit them to boxB sites located in the 3′ UTR of a constitutively expressed GFP reporter mRNA. To ensure that each cell expresses only one tagged ORF, we transduced reporter cells with retroviruses carrying ORFtag cassettes at low multiplicity of infection (MOI) followed by selection. Cells with altered GFP reporter expression – increased for activator and decreased for repressor and PTGR screens – were isolated by FACS and insertion sites were determined in pool by inverse PCR (iPCR) on ring-ligated short DNA fragments obtained by restriction digest and subsequent NGS^11^ (**Fig. 1a**). Finally, we identified gene loci where insertions were statistically over-represented in the sorted samples compared to the non-selected background dataset by assigning each integration to the nearest downstream splice acceptor-containing exons of protein-coding genes (see **Online Methods** and **Extended Data Table 1**). For each of the three screens, we found a prominent, screen-specific enrichment of insertions at positive control genes, exemplified by the transcriptional coactivator Yap1 (for the activator screen), the KRAB domain-containing Zfp57 (repressor), and the mRNA deadenylase complex subunit Cnot9 (PTGR) (**Fig. 1c**).

Overall, we identified 139 putative transcriptional activators, 207 repressors, and 77 PTGR proteins using stringent thresholds (FDR < 0.1%, log2 odds ratio ζ1, see **Extended Data Table 1**). Activator hits include several known transcriptional activators, such as p65, Ep300, subunits of the Mediator complex, and all Kmt2(a-d) histone-methyltransferases, which could not be screened before due to their long ORFs of up to 17kb (**Fig. 1d**, **Extended Data Fig. 1a**). Repressor hits contain 75 KRAB zinc-finger repressors and their corepressor Trim28, HP1 family proteins, H3K9 methyl-transferases, and Polycomb repressive complex components (**Fig. 1d**, **Extended Data Fig. 1a**). Finally, the PTGR screen identified core components of the microRNA (Ago2, Tnrc6a/b/c) and nonsense-mediated decay (Smg1, Smg9, Upf2) pathways, members of the Ccr4-Not deadenylation complex (Cnot2, Cnot3, Cnot9), as well as translational inhibitors (Eif4e2, Eif4enif1) (**Fig. 1d**, **Extended Data Fig. 1a**).

While ORFtag integrations per screen were highly reproducible (**Extended Data Fig. 1b**), the hits from the three distinct assays showed almost no overlap, indicating that ORFtag does not lead to the recurrent and artefactual detection of unspecific genes (**Fig. 2a & Extended Data Fig. 1c**). Consistent with this, the activator and repressor screen hits were strongly enriched for proteins containing activating and repressive domains, respectively, and both protein sets share a significant enrichment for known transcription factors (TFs) (**Fig. 2b**). Similarly, only PTGR hits were enriched for known RNA-binding proteins (**Fig. 2b**). Moreover, the genes identified by the three screens were enriched for distinct gene ontology (GO) terms, protein domains and subcellular locations, all of which are consistent with their associated functions (**Fig. 2c**). Overall, these results indicate that ORFtag is compatible with diverse functional assays and delivers assay-specific hits.

**Fig. 2:**
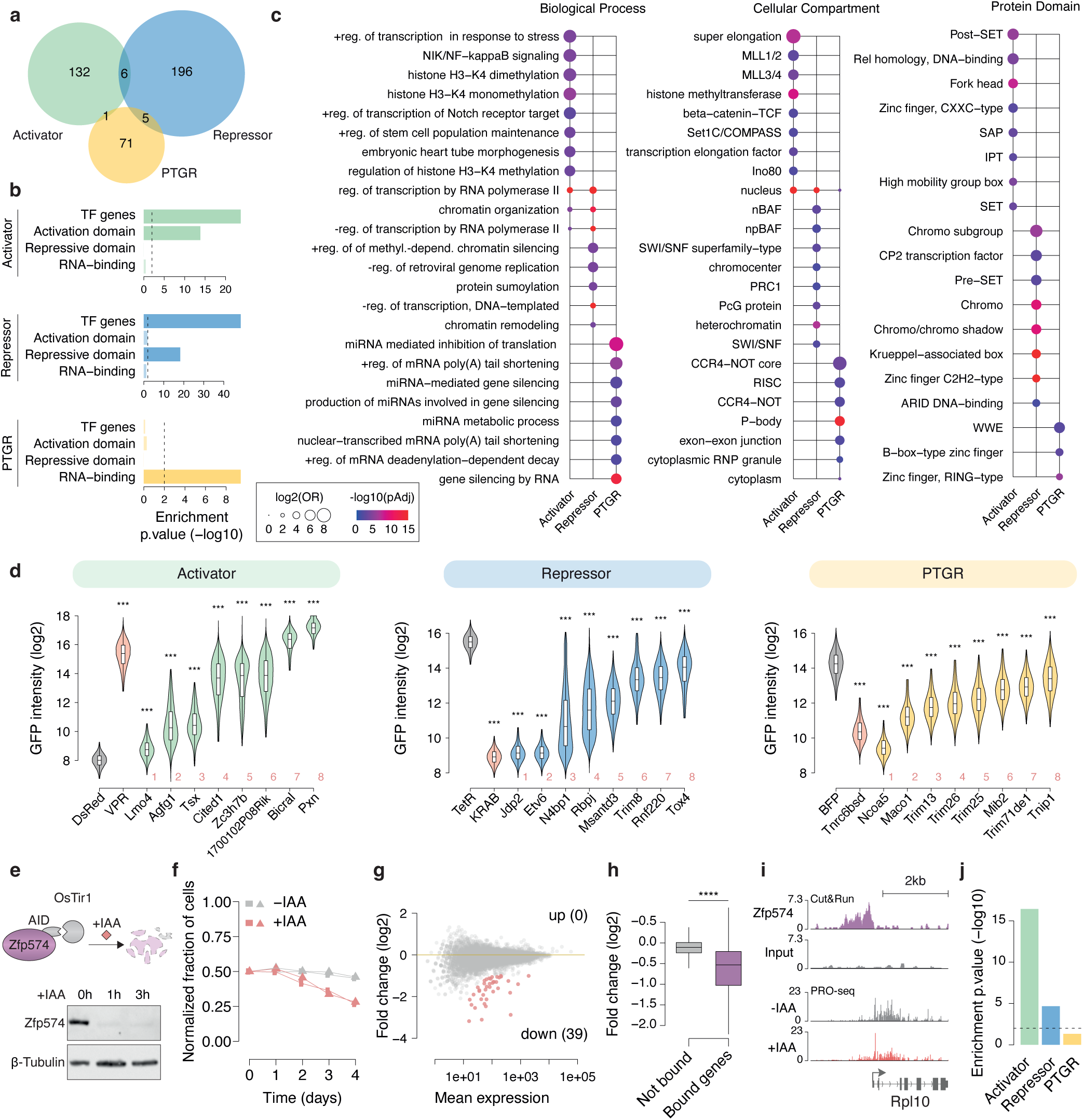
ORFtag interrogates protein function with high specificity. **a,** Overlap between activator (green), repressor (blue) and PTGR (yellow) hits. **b,** Enrichment of screen hits for human homologous genes with annotated DNA-binding, activation, or repressive domains and RNA-binding proteins. **c,** Top enriched protein domains, biological process and cellular compartment GO terms for activator, repressor and PTGR hits. **d,** Independent validation of select screen hits. GFP intensity measured by flow cytometry in reporter cell lines stably expressing the indicated full-length proteins fused to TetR (Activator, Repressor) or λN (PTGR); Wilcoxon test, ***p≤1×10^−3^. Refer to Fig. 1d for the position of the hits in the volcano plot. **e**, Schematic view of Zfp574 rapid depletion using Auxin-Inducible Degron (AID). The rapid depletion of Zfp574 upon 3-indoleacetic acid (IAA) treatment shown by Western Blot. **f**, Cell viability timecourse in the presence (-IAA, in grey) or absence of Zfp574 (+IAA, in red). Shown are two biological replicates. **g**, MA plot showing PRO-seq fold-changes (log2) upon Zfp574 6h depletion. Signficantly up- (0) or down-regulated (39) genes are highlighted in red. **h**, PRO-seq fold-changes (log2) of not-bound versus Zfp574 promoter-bound genes upon Zfp574 depletion; Wilcoxon test, ****p≤1×10^−5^. **i**, Zfp574 Cut&Run and PRO-seq screenshots at the Rpl10 locus. **j**, Enrichment of screen hits for genes that were identified as part of oncogenic fusions.

To experimentally validate the screen results at the level of protein-inherent functionality, we individually cloned and transduced eight hits from each screen, fused to the respective TetR or *λ*N tags, and tested whether they were sufficient to regulate the respective reporters. All candidates tested, including hits that were not previously linked to the respective biological processes, could be validated in recruitment assays together with previously known regulators, confirming that ORFtag screens are highly specific and have low false positive rates (**Fig. 2d**). For example, recruitment of the annotated cytoskeletal protein Pxn or the uncharacterized protein 1700102P08Rik was sufficient to strongly activate transcription, while the E3 ubiquitin ligase Trim8 and the uncharacterized protein Msantd3 were sufficient to repress transcription. In turn, the neuronal activity-associated protein Maco1 and the E3 ubiquitin ligase Trim13 were sufficient to repress reporter gene expression when recruited to the 3’UTR of an mRNA.

To assess the potential of ORFtag in assigning cellular roles to uncharacterized proteins, we sought to investigate the endogenous function of the zinc-finger protein Zfp574, which ORFtag specifically identified as a transcriptional activator. Utilizing the auxin-inducible degron system (**Fig. 2e**), we show that depletion of Zfp574 results in a significant growth defect (**Fig. 2f**), indicating that Zfp574 is essential for cellular fitness. Rapid depletion of Zfp574 followed by PRO-seq further revealed that Zfp574 functions strictly as a transcriptional activator in accordance with the ORFtag results (39 genes go down, 0 genes go up upon depleting Zfp574 at FDR≤0.05 and FC≥2) (**Fig. 2g**). Cut&Run for Zfp574 identified 140 binding sites genome-wide, the majority (87.9%) of which are located in promoter-proximal positions (+-500 bp around the gene transcription start sites), and transcription of the promoter-bound genes was strongly affected upon Zfp574 depletion (**Fig. 2h, i**). Thus, Zfp574 is a novel selective transcriptional activator that specifically binds and activates a small set of genes that support cell fitness. Taken together, our results demonstrate that ORFtag, coupled with functional assays, provides a robust and powerful method for the high-throughput assignment of protein function.

Some identified hits may regulate gene expression in ORFtag assays without necessarily doing so endogenously. This underscores the distinction between a protein’s inherent biochemical function (as evaluated here) and its role within the cell. In fact, the process of tagging and/or chromatin- or RNA-tethering can potentially overwrite a protein’s usual cellular function and change its localization within the cell (e.g. signaling peptides can be bypassed, replaced or overwritten by ORFtag). Importantly, these hits remain valuable as their ability to activate/repress gene expression *in principle* is highly relevant, e.g. in cancer, when chromosomal re-arrangements create oncogenic fusion proteins. Indeed, among our hits is the ortholog of the oncogene C3orf62, recently described by a tethering-based approach to be an activator^2^. We compared the ORFtag hits systematically to their human orthologs and found oncogenes to be enriched among the activators and – more weakly – the repressors but not the post-transcriptional regulators (**Fig. 2j**). These include for example Zc3h7b and D630045J12Rik (KIAA1549in human) that can function as activators, and Gm10324 that can function as a repressor, highlighting that oncogenic fusions can recruit unrelated genes to function in gene regulation.

Having established that retroviral integration sites represent indeed successful ORF tagging events that score in functional assays, we conducted a systematic and critical assessment of ORFtag’s ability to comprehensively and reproducibly tag proteins. Comparison of the retroviral integration sites from six independent transductions, performed in three different laboratories, revealed that, regardless of the protein tag used, experimental cassettes integrated in a similar distribution across the genome and the number of insertions per genomic region correlated highly (PCC≥0.84) (**Extended Data Fig. 1b**). ORFtag integrations were enriched near transcription start sites (TSS), a well-known feature of retroviral vectors^11^, enabling the tagging of near-full-length proteins (**Fig. 3a**). Assigning each integration to a gene locus indicated that we were able to tag at least 83.7% of all mouse protein-coding genes with a median count of 15 integrations per gene, given the scale and sequencing depth of our screens (**Fig. 3b, 3c**). The tagged genes include those with large open reading frames yielding high molecular weight proteins (**Fig. 3d**). Indeed, in contrast to ORFeome-based approaches that are biased towards short ORFs, ORFtag is not influenced by gene length (**Fig. 3e**). Moreover, the retroviral ORFtag cassette allowed the tagging of ORFs that exhibited different endogenous expression levels, including >59% of genes that are normally not expressed in mESCs (**Fig. 3f**). Importantly, also the hits identified in the three functional screens include genes of varying lengths and expression levels (**Fig. 3e & 3f**).

**Fig. 3:**
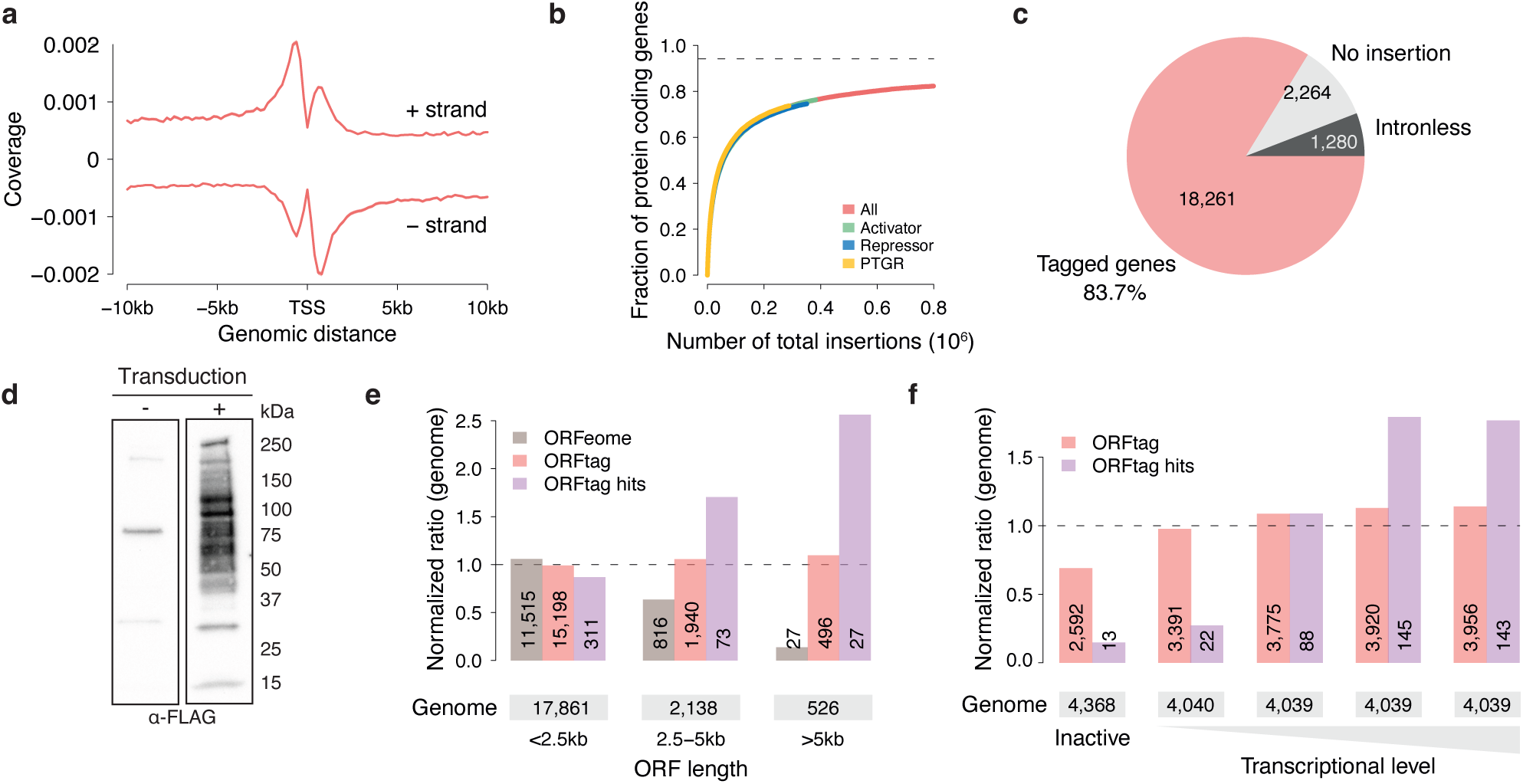
Scope and limitations of massive parallel protein tagging using ORFtag. **a,** Distribution of ORFtag integrations around TSSs of mouse protein-coding genes. **b,** Saturation curve displaying the relationship between the fraction of tagged proteins and the number of determined integration sites. **c,** Fraction of genes showing at least one integration in the combined background sample. **d,** Western blot against the FLAG tag assessing the tagging pattern in mESC lysate before (-) and after (+) ORFtag transduction. **e,** Ratio of protein coding genes that were successfully tagged using ORFtag (ORFtag, pink) or were hits in any of the three screens (ORFtag hits, purple), normalized by the distribution found across the whole mouse genome (Genome, dashed line). Human ORFeome is shown for comparison (ORFeome, light grey). See material and methods for further details. **f,** Ratio of intron-containing protein coding genes that were successfully tagged using ORFtag (ORFtag, pink) or were hits in any of the three screens (ORFtag hits, purple), normalized by the distribution found across the whole mouse genome (Genome, dashed line).

A limitation of ORFtag lies in its inability to functionally probe intronless genes and first exons, due to them lacking splice acceptor sites. However, it is worth noting that 45.6% of first exons are non-coding and that among protein-coding first exons, the median encoded peptide length is 31 amino acids short. As a result, only 12.8% of first exons contain annotated protein domains (**Extended Data Fig. 1d**). Intronless genes, which cannot be tagged, constitute only a small fraction of protein coding genes (5.9%). These are dominated by a few protein families, including histones and various sensory receptors (**Extended Data Fig. 1e**), leaving 94.1% of protein-coding genes as potentially taggable by ORFtag. We also note that certain genes may not be accessible to ORFtag screens if cellular fitness is sensitive to changes in their expression levels.

In conclusion, ORFtag is an easy-to-implement functional genomics tool that enables cost-effective proteome-scale functional screens. Due to its modularity, ORFtag can be combined with various functional tags and a wide range of applications, including bio-imaging, targeting proteins to various organelles, and protein-protein interaction studies. Notably, while ORFtag utilizes N-terminal tagging, it can be adapted for C-terminal or internal tagging, broadening the scope of proteins and applications that can be explored. These alternative tagging approaches would mirror the tagged genes’ endogenous expression levels and are thus limited to genes expressed in the particular cell line. Finally, ORFtag can be readily employed in cellular systems of various model organisms without the need to generate species-specific resources. This adaptability and versatility make ORFtag a promising tool for advancing functional genomics research.

## Extended Data Legends

**Extended Data Fig. 1:**
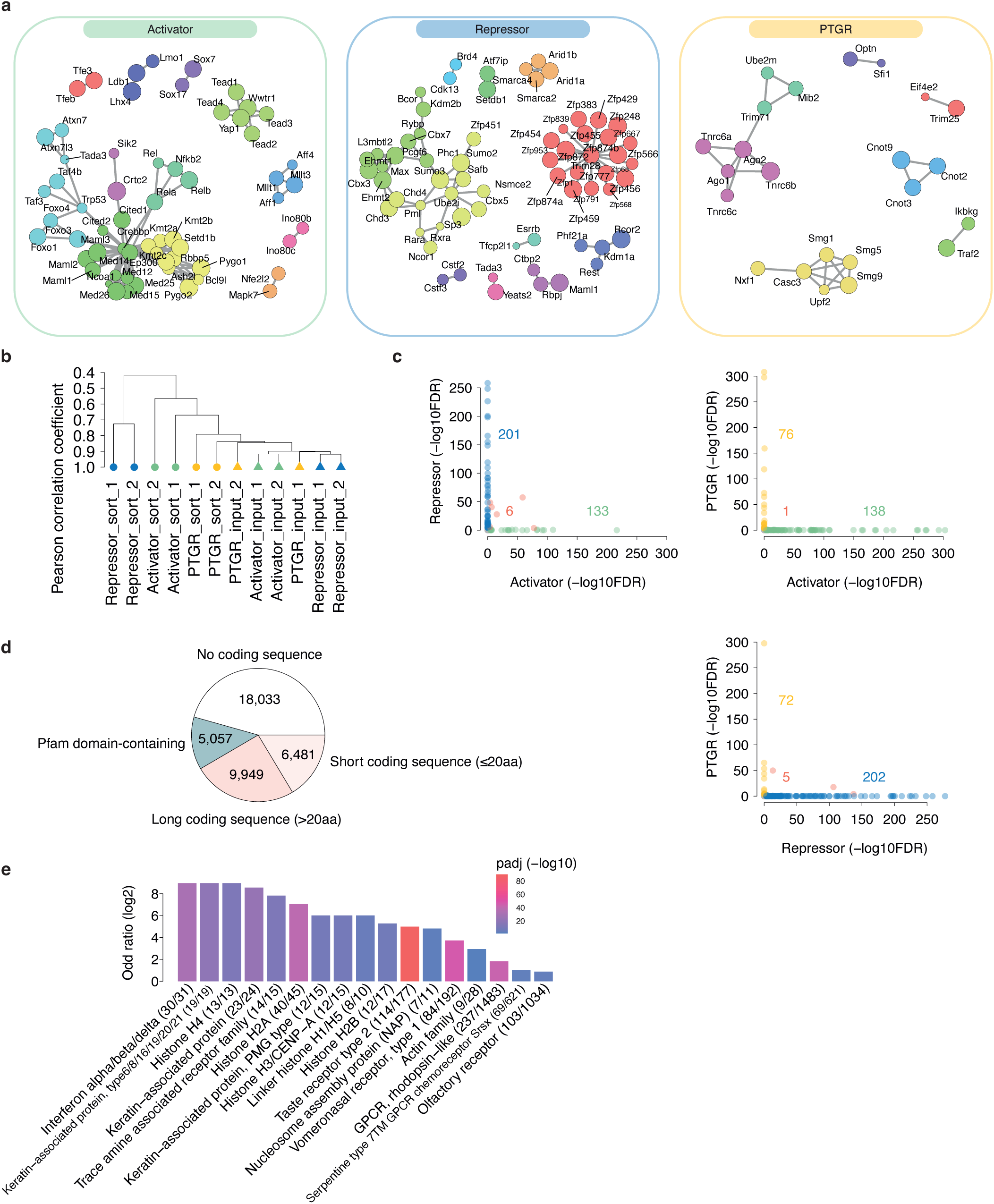
Evaluation of ORFtag integrations at global scale. **a,** STRING protein-protein interaction networks between activator/ repressor/ PTGR hits. Node communities were highlighted using a color code (Louvain method), and their size is proportional to the Odd ratio of the corresponding hit. Only the hits showing at least one interaction with another hit are shown. **b,** Dendrogram of Pearson’s Correlation Coefficients between unsorted (triangles) and sorted (round) replicates from each functional screen. All background (input) samples show high PCC (≥ 0.85). **c,** Scatter plots displaying a pairwise comparison of −log10(FDR) values for screened genes across different assays. **d,** Protein family enrichment of intronless genes, whose majority belongs to few protein families. **e,** Fraction of exons that are non-coding (in white), code either for no known protein domain (in shades of pink) or for a high confidence PFAM protein domain (Domain-containing CDS, in blue). Importantly, most first exons are non-coding or do not contain specific protein domains, fostering the use of ORFtag for a wide range of functional studies.

**Extended Data Table 1: Identification of activator, repressor and PTGR hits**

For each gene locus, raw counts, odd ratio (log2) and the associated FDR are shown for the 3 different screens. The last “hit” column specifies whether a locus was considered as a hit (TRUE) or not (FALSE).

## Methods

### Cell culture conditions

All experiments presented here were carried out in diploid mouse embryonic stem cells (mESCs) that were derived from originally haploid HMSc2 termed AN3-12^11^. The mESCs were cultivated without feeders in high-glucose-DMEM (Sigma-Aldrich) supplemented with 13.5% fetal bovine serum (Sigma-Aldrich), 2 mM L-glutamine (Sigma-Aldrich), 1x Penicillin-Streptomycin (Sigma-Aldrich), 1x MEM non-essential amino acid solution (Gibco), 1mM sodium pyruvate (Sigma-Aldrich), 50 mM β-mercaptoethanol (Merck) and in-house produced recombinant LIF. Virus packaging cell lines, Lenti-X 293T (Takara), and PlatinumE (Cell Biolabs), were grown according to the manufacturer’s instructions. All cell lines were cultured at 37°C and 5% CO2 and regularly tested for mycoplasma contamination.

### Reporter cell lines

The reporter cell line for the Repressor screen was established previously^12^ and contains the reporter construct inserted into the expression-stable locus on Chr15 that is compatible with the Flp recombinase-mediated cassette exchange (RMCE). Reporter cell line for the Activator screen was generated by RMCE as follows – 5×10^6^ cells were electroporated with a mix of 10 µg of plasmid containing constructs flanked by FRT/F3 sites, and 6 µg of plasmid expressing Flp, using a Maxcyte STX electroporation device (GOC-1) and the Opt5 program. Seven days after the transfection, cells were sorted and clonal cell lines were generated. Cell lines were genotyped using integration-site specific PCRs and Sanger sequencing. The Activator reporter construct (Addgene, this study) contains the PuroR-IRES-GFP reporter under the control of the minimal promoter derived from the MYLPF gene (chr16:30374730−30374857 +, hg38) that was shown to have a low basal expression and high inducibility^13^. Upstream of the promoter are 7x TetO sites flanked by the loxP sites.

The reporter cell line for the PTGR screen was created by nucleofection of haploid AN3-12 mESCs with 500ng of the reporter construct and 10 µg of a Tol2 transposase encoding plasmid using the Mouse ES Cell Nucleofector Kit (Lonza) according to the manufacturer’s protocol using an Amaxa Nucleofector (Lonza). The PTGR reporter construct (Addgene, this study) encodes for PuroR-IRES-GFP followed by 10 boxB sites that are flanked by two loxP sites under the control of a PGK promoter. Cells were subsequently selected using 1 µg/ml Puromycin (Gibco) followed by single clone selection. Single cell clones were afterwards transduced with a retroviral vector for the expression of pMSCV_hygro_CreERT2 and selected with 250 µg/ml Hygromycin (Roche) followed by single cell clone selection.

### ORFtag screens

The ORFtag viral constructs were derived from the ecotropic Retro-EGT construct^11^ that includes the sequence features necessary for the inverse-PCR protocol (see below). Furthermore, the construct contains constitutively active PGK promoter that drives the expression of a NeoR resistance gene separated from a tag by the IRES sequence. The tag contained either TetR with an N-terminally located nuclear localization signal (Activator and Repressor screens; this study) or LambdaN domain (PTGR screen; this study). Additionally, the tag contained 2x GGGS-linker followed by the BC2-tag and 3xFLAG-tag. Finally, the ORFtag construct contains a consensus splice donor motif followed by a part of the Hprt intron (chrX:53020400-53020556 +, mm10). In order to tag genes in all three possible coding frames, three constructs were used that contain either 0, 1 or 2 additional nucleotides upstream of the consensus splice motif.

Every ORFtag screen was performed in two independent replicates. Retroviral constructs carrying ORFtag cassette were packed in PlatinumE cell lines using polyethylenimine (PEI) reagent as described previously^11^. Reporter cell lines (100-150 million cells) were transduced with packaged retrovirus in the presence of 6 µg/ml polybrene (Sigma) and at low transduction efficiency (< 20%) to ensure only one virus per cells. Cells were harvested 24 hours later and plated in medium containing 0.1 mg/ml G418 (Gibco) form selection of transduced cells. Selection was continued until all cells on the control plate died (4-5 days), after which 40 million cells were processed as non-selected background for mapping of genomic integrations (see below). The remaining cells were sorted for GFP-positive (Activator screens) or GFP-negative (Repressor screens) populations using BD FACSAria III or IIu cell sorters (BD Biosciences) and processed for mapping of genomic integrations (see below). For the PTGR screen a five-sort strategy was applied to enrich cells that show a tethering dependent repression of reporter gene expression. Cells with a GFP expression equal to the lowest 10 percent of GFP expression observed after selection were sorted using BD FACSAria III and expanded thereafter. Additionally, non-sorted cells were maintained for gating of the consecutive sorts. Two additional sorts for cells with GFP expression similar to the lowest 10% of GFP signal observed in the non-sorted cells were performed and again expanded in-between the sorts. A fourth sort was performed for cells with a GFP expression equal to the lowest 5% of GFP signal observed in the non-sorted cells. After expansion, the cells were treated with 500nM 4-Hydroxytamoxifen (Sigma) to induce Cre-mediated recombination and to flox the boxB sites of the reporter construct and hence to revert the tethering. Thereafter a final sort was performed to select a cell population with a GFP expression equal to the highest 70% of GFP expressing cells.

Transduction efficiency was measured by plating 10,000 cells on a 15-cm dish and selecting with G418 (Gibco). A control plate with 1,000 cells was also plated without selection. After 10 days, colonies were counted and transduction efficiency was calculated as the number of colonies on the selected plate divided by the total number of cells plated (10 times the number of colonies on the control plate).

### Mapping of genomic integrations by next-generation sequencing

Genomic locations of ORFtag integrations were mapped using modified inverse-PCR followed by next generation sequencing (iPCR-NGS) protocol^11^. Genomic DNA was prepared by lysing cell pellets in lysis buffer (10 mM Tris-HCl pH 8.0, 5 mM EDTA, 100 mM NaCl, 1% SDS, 0.5 mg/ml proteinase K) at 55°C overnight. Following a 2-hour RNase A treatment (Qiagen, 100 mg/ml, 1:1,000 dilution) at 37°C, two extractions using phenol:chloroform:isoamyl alcohol and one extraction using chloroform:isoamyl alcohol were carried out. The samples then underwent two separate digestion reactions (with up to 4 µg of genomic DNA) using NlaIII and MseI enzymes (NEB) at 37°C overnight, followed by purification using a Monarch PCR&DNA Cleanup Kit (NEB). Ring-ligation was carried out using T4 DNA ligase (NEB) at 16 °C overnight, followed by heat-inactivation (65°C, 15 min) and linearization using SbfI-HF (NEB) at 37°C for 2 h. The digests were then purified using a Monarch PCR&DNA Cleanup Kit (NEB) and amplified using firstly a nested PCR reaction with KAPA HiFi HotStart ReadyMix (Roche), and a specific primer pair (TGCAGGACCGGACGTGACTGGAGTTC*A, TGCAGGACGATGAGCAGAGCCAGAACC*A) for 16 cycles. After cleanup with AMPure XP Reagent (Beckman Coulter, 1:1 ratio beads:PCR), iPCR amplification was carried out with KAPA HiFi HotStart ReadyMix (Roche), and a specific primer pair (AATGATACGGCGACCACCGAGATCTACACGAGCCAGAACCAGAAGGAACTTGA*C, CAAGCAGAAGACGGCATACGAGAT [custom-barcode] GTGACTGGAGTTCAGACGTGTGCTCTTCCGATCT) for 18 cycles. Afterwards, amplified libraries were size selected for a range of 400-800 bp using SPRIselect beads (Beckman Coulter). NGS was performed on an Illumina NextSeq550 or llumina HiSeq 2500 sequencer according to the manufacturers’ protocols with custom first-read primer (1:1 mix of GAGTGATTGACTACCCGTCAGCGGGGGTCTTTCA and TGAGTGATTGACTACCCACGACGGGGGTCTTTCA).

### Immunoprecipitation

To confirm expression of tagged proteins, the PTGR reporter mESCs, transduced with the ORFtag construct as well as non-transduced cells, were lysed in lysis buffer (50mM TRIS HCl pH: 7.5, 150mM NaCl, 0.1% SDS, 1% Triton-X-100, 0.5% NP-40, 0.5mM EDTA supplemented with Proteinase Inhibitor (Roche) and protein concentration was determined photometrically using the Protein Assay Dye Reagent Concentrate (BioRad), according to the manufacturer’s protocol and photometric measurement at 595nm. Tagged proteins were captured using 80 µl of in-house BC2-nanobody coupled magnetic beads from 1mg total protein. Bound proteins were eluted by resuspension of the beads in 1x SDS-sample buffer and incubated at 95°C for 5 minutes. Further details about Western blotting can be found below.

### Individual recruitment validations

To validate Activator hits, the candidates were amplified by PCR from mESC cDNA and inserted into retroviral constructs that comprises the PGK promoter that drives the expression of a PuroR resistance gene and a tag separated by the IRES sequence. The tag contains TetR, along with an N-terminal nuclear localization signal, a 2x GGGS-linker, a BC2-tag, and a 3xFLAG-tag, followed by the tested candidate. Retroviral constructs were packed in PlatinumE cell lines (see above), and reporter cell lines (170,000 cells) were transduced in the presence of 6 µg/ml polybrene (Sigma). Cells were harvested 24 hours later and plated in medium containing 1 µg/ml Puromycin (InvivoGen) to select for transduced cells. After five days of selection, the reporter expression was analyzed on an LSR Fortessa (BD) flow cytometer. For processing and visualization, FlowJo and R package flowCore (v2.12.2) was used.

In order to validate Repressor and PTGR hits, PCR was used to amplify the candidates from mESC cDNA, and lentiviral plasmids were created as fusion proteins containing TetR/lamdaN-Candidate-P2A-mCherry coding sequence under the control of an EF1a promoter. For the validation of Trim71, cDNA excluding the fragment encoded in exon 1 (Trim71dE1) was cloned into the aforementioned lentiviral plasmid. A fragment encoding for the silencing domain of human Tnrc6b (Tnrc6b-SD) was expressed using the same lentiviral plasmid as above as a positive control for the validation of PTGR hits. Lentivirus was produced in Lenti-X 293T cells as in (Ref.^12^). Reporter cells were then transduced with the virus in the presence of 8 µg/ml polybrene (Santa Cruz Biotechnology, SACSC-134220). After 7 days of transduction, reporter expression was analyzed on an LSR Fortessa (BD) flow cytometer. Reporter cells transduced with recruitment constructs were gated based mCherry expression. For processing and visualization, FlowJo and R package flowCore (v2.12.2) was used.

### AID cell line generation

A parental cell line expressing the E3 ligase for AID was generated by inserting a cassette into the expression-stable locus on Chr15 that is compatible with the Flp recombinase-mediated cassette exchange in mESCs (RMCE, see “Reporter cell lines” section). The construct contained EF1alpha-ARF16-HA-P2A-OsTir1-3xMyc-T2A-mCherry-SV40_polA site flanked by the FRT/F3 sites. The clonal Tir1 parental cell line was genotyped using integration-site specific PCRs and Sanger sequencing.

To generate the N-terminally AID-tagged Zfp574 cell line, 5×10^6^ Tir1 parental cells were transfected with 10 µg of plasmid (Ref.^14^) that expresses Cas9 and the gRNA against a target locus (CTTGCTGCTGCCATGACTG) and 5 µg of plasmid with a knock-in cassette containing Blasticidin-P2A-V5-AID-GGGS flanked by 20 bp microhomology arms (Ref.^14^) using a Maxcyte STX electroporation device (GOC-1) and the Opt5 program. Two days after the transfection, cells were selected for knock-ins with 10 µg/mL Blasticidin (ThermoFisher), individual clones were genotyped using knock-in-site specific PCRs and Sanger sequencing. Potential candidates were investigated by western blotting against the integrated V5-tag (Thermo Fisher, R960-25) with or without 500 uM 3-indoleacetic acid (IAA, Merc) treatment.

### Western blotting

Cells (3×10^6^) were collected, centrifuged at 300g for 5 min, washed with 1xPBS and lysed in 100 µl RIPA buffer containing protease inhibitor (Roche) and Benzonase (Sigma Aldrich). For complete lysis, cells were incubated on ice for 30 min and sonicated for five minutes (30 sec on/off, Diagenode Bioruptor). Afterwards, samples were centrifugated for 5 min at full speed and 4 °C, and supernatants were supplemented with 20 µl 4x Laemmli buffer with 10% β-mercaptoethanol. Samples were boiled for 5 min at 98°C.

Proteins were resolved on SDS-PAGE on a 4-15% Mini-PROTEAN TGX gel (BioRad) and transferred to an Immobilon-P PVDF membrane (Merck Millipore) using a wet-chamber system. Tagged proteins were detected using mouse α-Flag M2 (Sigma Aldrich F3165, 1:10,000), mouse α-V5-tag (Thermo Fisher R960-25, 1:1,000), or rabbit α-β-tubulin (Addgene ab6046, 1:10,000) as primary and HRP-α-Mouse (Cell Signaling, 7076, 1:10,000) or HRP-α-Rabbit (Cell Signaling, 7074, 1:10,000) as secondary antibody and imaged using ClarityTM Western ECL Substrate (BioRad) with a ChemiDocTM Imaging System (BioRad) using ImageLab v5.1.1 (BioRad).

### Cell viability timecourse

For growth curve assays, AID-tagged cell line (mCherry positive, see AID cell line generation) was mixed at 1:1 ratio with WT cells, split into control (-IAA) and treatment (+IAA, Merc, 500 uM) group and cultured in a 24-well cell culture plate. The ratio between mCherry positive and negative cell was quantified every 24hrs by Flow Cytometry (iQue Screener PLUS, Intellicyt).

### PRO-seq

For each condition, 1×10^7^ AID-Zfp574 cells were collected and nuclei were isolated after 6h of 500uM IAA treatment or no treatment (two biological replicates per condition). Spike-in control (S2 *Drosophila* cells; 1% of mESC cells) were added at the level of nuclei permeabilization step. The next steps of the PRO-seq protocol were performed as in (Ref.^15^) with a single modification: the nuclear run-on was performed at 37 °C for 3 min.

### Cut&Run

For each biological replicate, 1×10^6^ cells from the AID-Zfp574 cell line or the Tir1 parental cell line were used. The Tir1 parental cell line is used as Input, each experiment was performed in two biological replicates. The protocol was performed as in (Ref.^16^) with a V5-tag antibody (Thermo Fisher, R960-25) that was added to a final dilution of 1:100.

### Bioinformatic analyses

All bioinformatic analyses were performed in R (v4.2.0, https://www.R-project.org/). Computations on genomic coordinate were conducted using the GenomicRanges (v1.50.1)^17^ and the data.table (https://CRAN.R-project.org/package=data.table) R packages. All box plots depict the median (line), upper and lower quartiles (box) ±1.5x interquartile range (whiskers); outliers not shown.

### Processing of ORFtrap screens

First, iPCR reads from sorted and background (non-selected) samples were trimmed using Trim galore (v0.6.0) with default parameters to remove Illumina adapters. Then, trimmed reads were aligned to the mm10 version of the mouse genome using bowtie2^18^ with default parameters (for paired-end sequenced samples, only first mate reads were considered), before removal of duplicated and low mapping quality reads (mapq<=30) using samtools (v1.9)^19^. Mapped insertions were assigned to the closest downstream exon junction – with a maximum distance of 200kb – based on GENCODE annotations of the mouse genome (vM25). Finally, insertion counts were aggregated per gene. Of note, only exons from protein-coding transcripts were considered, except for the first exon of each transcript, which might not contain splicing acceptor sites. Consequently, intronless genes – for which none of the isoforms contain a spliced intron – were not considered.

Background replicates showed reproducible gene counts (PCC ≥0.84) and therefore were merged, and genes with at least one insertion were considered as putatively tagged. Finally, genes showing significantly more insertions in sorted samples compared to merged background samples were identified using one-tailed fisher’s exact test (alternative= “greater”) on merged biological replicates. Of note, only genes with at least 3 unique insertions in sorted samples were considered. Obtained p-values were corrected for multiple testing using the FDR method and genes showing an FDR<0.001 and a log2 Odd Ratio≥1 were classified as hits.

### Protein-protein interaction networks

For each functional assay, STRING protein-protein interaction between hits were retrieved using the STRINGdb R package (v2.10.0, database version 11.0). Finally, only the hits showing at least one protein-protein interaction with another hit with a combined score ≥900 were considered.

### CDS length bias

To assess whether ORFtag is biased towards short ORFs, we stratified intronic protein coding genes based on their shortest CDS length (< 2.5kb, 2.5-5kb and longer than 5kb). Then, we compared how tagged genes (with at least one insertion in background samples) and hits (union from the three screens) were distributed between these groups, using all intronic protein coding genes as a reference. For example, to compute the normalized ratio of tagged genes for the <2.5kb group, we used the following formula: normalized ratio= ([tagged genes with CDS<2.5kb]/[total tagged genes])/([intronic protein genes with CDS<2.5kb]/[total intronic protein coding genes]). To allow side-by-side comparison, we also considered ORFs from the human ORFeome that Alerasool and colleagues were able to transfect and detect^2^.

### Gene expression bias

To assess whether transcriptionally inactive mouse genes could be assayed using ORFtag, we used publicly available data from the same mESC cell line (GSE99971)^20^. For each intronic protein coding gene, mean TPM was computed across three RNA-Seq replicates (only protein-coding genes were considered). Genes with a mean TPM of 0 were classified as inactive and active genes were further stratified into quartiles. Then, we compared how tagged genes (with at least one insertion in background samples) and hits (union from the three screens) were distributed between these groups, using all intronic protein coding genes as a reference. For example, to compute the normalized ratio of tagged genes for the inactive group, we used the following formula: normalized ratio= ([tagged genes with TPM=0]/[total tagged genes])/([intronic protein genes with TPM=0]/[total intronic protein coding genes]).

### Enrichment analysis of expected protein functions

To assess whether hits were enriched for genes with expected functions, we collected publicaly available lists of human TF genes (Ref.^21^), human genes containing activation or repressive-domains (Ref.^22^, only genes containing sufficient (‘S’ or ‘N and S’) and high confidence (‘H’) domains were considered), human genes containing RNA-binding domains (RBPbase^23^, only the genes identified in at least two different cell lines were used) and human fusion oncoproteins (COSMIC database v97, Ref.^24^). Screen hits were first assigned to their human orthologs using MGI^25^ homology data. For each functional assay, we assessed whether relevant categories were enriched among the hits using one-tailed fisher’s exact test (alternative= “greater”), and the total number of intronic protein coding genes as background.

### GO terms and protein domains enrichment

Biological Process (BP), Molecular Process (MF) and Cellular Compartment (CC) Gene Ontology (GO) terms were obtained from the org.Mm.eg.db (v3.15.0) R package. Protein domains were retrieved from the EnsDb.Mmusculus.v79 R package (v2.99.0). For each functional assay, GO terms and protein domains that were over-represented among hits were identified using one-tailed fisher’s exact test (alternative= “greater”), using all intronic protein coding genes as background. Obtained p-values were corrected for multiple testing using the FDR method and features with an FDR<0.05 were considered as significantly enriched. Of note, small categories containing less than five genes in total and categories with less than three matching hits were not considered. Finally, top 8-10 enriched GO terms and proteins domains were plotted for each functional assay.

### Protein family enrichment

To identify protein families enriched among intronless genes (for which none of the isoforms contain a spliced intron), annotations were retrieved from the EnsDb.Mmusculus.v79 R package (v2.99.0). Enriched protein families were identified using one-tailed fisher’s exact test (alternative= “greater”), and the total number of protein coding genes as background. Obtained p-values were corrected for multiple testing using the FDR method, and the protein families with an FDR<0.05 were plotted.

### Analysis of first exons

For the analysis of first exons, first exons containing a predicted CDS were classified as either short (≤20aa) or long (>20aa). Then, manually-curated Pfam-A domains from UCSC^26^ were used to discriminate first exon CDSs containing a know protein domain (e.g. coding for at least 10% of a full Pfam domain) or not.

### Gene annotation for PRO-seq analysis

To obtain a non-redundant set of genes for quantification of PRO-seq signals, we collected all coding and long non-coding transcripts from Ensembl v.100 for the mm10 version of the mouse genome, excluding transcripts shorter than 300 bp. When several transcript isoforms shared the same annotated transcription start site (TSS), only the longest isoform was retained. Next, TSS positions were corrected using FANTOM5^27^ CAGE TSS clusters: for each unique annotated TSS, we identified the strongest CAGE TSS within a 1kb window centered on the annotated TSS, excluding the coding sequence. Finally, for each CAGE TSS, only the full length of the nearest transcript was used to count overlapping reads (see next section).

### PRO-seq analysis

PRO-seq libraries were sequenced in paired-end mode with 36-bp read length. To eliminate PCR duplicates, an 8-bp long unique molecular identifier (UMI) was incorporated at the 5′ end of the reads during the sample processing. Before mapping, the UMI was separated, and the Illumina adapters were trimmed using cutadapt v.1.18. Only reads with a length greater than 10 bp were then mapped using Bowtie v.1.2.2 ^28^, initially to the mm10 version of the mouse genome. The mapping allowed for up to 2 mismatches and reported only the best alignment (-m 1 --best --strata) for each read. To ensure the counting of unique nascent RNA molecules, reads that mapped to the same genomic location were collapsed based on their UMIs, allowing for up to 1 mismatch. To create the PRO-seq coverage signal with the exact positions of RNA pol II molecules, only the first nucleotide of each read (i.e. the 3’ end of nascent transcripts) was considered and the strand swapped to match the transcription direction. A non-redundant CAGE-corrected gene set was used to count the number of UMI-collapesed 1nt-long mapped PRO-seq reads that overlap them (see the “Gene annotation for PRO-seq analysis” section). Differential analysis was performed using DESeq2^29^ (v.1.22.2) and significantly up- or down-regulated genes were selected using FDR<0.05, log2 Fold Change ≥1 threshold.

### Cut&Run analysis

Single-end 50-bp long reads were mapped to the mm10 genome using bowtie v.0.12.9, allowing up to 3 mismatches and only uniquely mapping reads were retained. Afterwards, peaks were called for each individual replicate, as well as for the combined replicates against their respective input, using Macs2 v.2.1.2.1, with following settings: -f BEDPE - g mm -B --nomodel --extsize 300 --SPMR. The Macs2 generated BedGraph files that contain normalized coverage were converted into BigWig using bedGraphToBigWig. Given the high correlation between two replicates (PCC of 0.613 at a common set of peaks), only the merged sample was used for assigning bound genes if the peak was localized within +-500 bp around the gene transcription start sites.

## Data availability

The raw sequencing data are available from GEO (https://www.ncbi.nlm.nih.gov/geo/) under accession number GSE225972.

## Code availability

All custom scripts that were generated for this study were made publicly available at https://github.com/vloubiere/ORFtag_2023.

## Acknowledgements

We thank the IMP/IMBA/GMI and Max Perutz Labs core facilities for support, especially Flow Cytometry teams for provided outstanding support. Filip Nemčko was supported by a Boehringer Ingelheim Fonds PhD fellowship. Vincent Loubiere was supported by HFSP (LT000926/2020) and EMBO (790-2019) postdoctoral fellowships. Research in the Ameres group is supported by the European Union (ERC, RiboTrace, CoG-866166) and the Austrian Science Fund (FWF, 10.55776/F80). Research in the Stark group is supported by the Austrian Science Fund (FWF, 10.55776/P29613, 10.55776/P33157 and 10.55776/P36971). Basic research at the IMP is supported by Boehringer Ingelheim GmbH and the Austrian Research Promotion Agency (FFG, FO999902549). Research in Brennecke group is funded by the Austrian Academy of Sciences and the European Community (grant no. ERC-2015-CoG-682181). Next-generation sequencing was done at the Vienna Biocenter Core Facilities GmbH (VBCF) Next-Generation Sequencing Unit. For the purpose of Open Access, the authors have applied a CC BY public copyright license to any Author Accepted Manuscript (AAM) version arising from this submission.

## Author contributions

F.N. and M.H. implemented the ORFtag method and protocols. F.N. performed the activator screen, M.H. the PTGR screen, and R.Y. and V.L. the repressor screen. F.N., M.H. and R.Y. performed candidate validation experiments. F.N. performed all Zfp574 follow-up experiments. V.L., F.N. and A.S. developed the bioinformatic pipeline. V.L. and F.N. performed NGS data and downstream analyses. N.F. and M.P. helped with the experiments. U.E. and S.L.A conceptualized the ORFtag approach. S.L.A., A.S., U.E. and J.B. coordinated and supervised the work. All authors wrote the manuscript.

